# Influenza A Virus Infection Impairs Neuronal Activity in Human iPSC-Derived NGN2 Neural Co-Cultures

**DOI:** 10.1101/2025.08.26.672266

**Authors:** Feline F. W. Benavides, Annabel L. V. Kempff, Hilde Smeenk, Bas Lendemeijer, Marla Lavrijsen, Johan A. Slotman, Steven A. Kushner, Femke M. S. de Vrij, Lisa Bauer, Debby van Riel

## Abstract

Influenza A virus (IAV) infection is associated with a wide variety of neurological complications, of which mild complications like impaired cognitive functioning are most prominent. Even though several studies have shown that many influenza viruses can enter the CNS, the neuropathogenesis of seasonal (H3N2 and H1N1) and pandemic (pH1N1 2009) IAV infections is poorly understood. Therefore, we aimed to investigate the cellular tropism, replication efficiency and associated functional consequences using a human stem cell-derived neural co-culture model of neurons and astrocytes. All viruses were able to infect neurons in the co-culture model, although this infection did not result in efficient replication and release of progeny virus. In addition, infection did not result in visible cell death or apoptosis. However, functional analyses revealed that IAV inoculation resulted in a reduction of spontaneous neural activity and a partial reduction of neural excitability. This study shows that seasonal and pandemic IAVs can disrupt neural homeostasis, without efficient virus replication or the induction of cell death. However, these functional changes in neural activity can contribute to cognitive problems during IAV infections in the acute and potentially post-acute phase of the infection.

## Introduction

Influenza A virus infection causes tremendous global morbidity and mortality. Even though it is considered a respiratory pathogen, neurological symptoms can occur ^1–3^. Neurological complications vary in symptoms and onset and differ among influenza A virus subtypes. Zoonotic viruses, like H5Nx viruses, are particularly well known for their ability to cause severe neurological complications in a wide range of mammals, including humans^4^. Seasonal and pandemic influenza A viruses generally cause milder neurological complications, such as cognitive problems, and rarely more severe complications^5,6^.

There is evidence from patients and *in vivo* studies that seasonal and pandemic influenza A viruses can enter the CNS. In humans, both viral RNA and antigen have been detected in the brain post-mortem, and occasionally in cerebrospinal fluid^3,7–11^. In ferrets, infectious virus or viral RNA has been detected in the olfactory bulb or brains of seasonal H3N2 virus inoculated ferrets^5,12–18^ and in the olfactory bulb, cerebrum and cerebellum of pandemic H1N1 2009 (pH1N1 2009) virus inoculated ferrets^5,19^. In pandemic 1918 H1N1 inoculated ferrets, infectious virus, viral RNA and viral antigen were detected in the olfactory bulb, cerebrum, brain stem and trigeminal nerve^20^. In mouse studies, neuroinvasion has been observed with mouse-adapted H1N1 viruses (H1N1 WSN) and H3N2 virus^21,22^, while the pandemic H1N1 2009 virus (pH1N1 2009) and mouse-adapted H1N1 (H1N1 PR8) could not be detected in the brain^21,23,24^. Even though neuroinvasion is observed for a variety of seasonal and pandemic influenza viruses, the neurotropism has not been studied extensively. Previous *in vitro* studies have shown that pH1N1 2009, seasonal H3N2, H1N1 WSN and H1N1 PR8 can infect and replicate in neural cells^22,25–27^. However, these studies use either cell lines, lab-adapted or mouse-adapted viruses and have not been performed in more relevant human neural culture systems and/or with relevant clinical isolates.

Knowledge of the neurovirulent potential of seasonal and pandemic influenza A viruses is limited. For seasonal viruses, mild neurological complications, like learning and memory problems are most prominent^28^, but severe neurological complications like encephalitis, transverse myelitis, brain edema or loss of vision have been reported^29–36^. For pandemic viruses, like during the 2009 H1N1 pandemic, severe neurological symptoms were more frequently reported and included seizures and encephalopathies^37–39^, but also milder symptoms like confusion, dizziness and headache^2,36,39–41^. Additionally, a recent study revealed that (severe) influenza A virus infections increase the risk for neurodegenerative diseases^42^. Even though neurological complications associated with influenza A virus infections have been described since 1918^43^, comprehensive knowledge about the neuropathogenesis, and especially that of less severe complications, such as memory and learning problems, is still lacking.

Neurodegenerative and neuropsychiatric diseases, especially those with learning and memory problems, are associated with neuronal cell death, decreased neural activity, and impaired synaptic plasticity^44,45^. Influenza A virus infection could lead to cell death by the cytopathic effect of the infection or through the direct or indirect induction of apoptosis. An increase of the apoptotic marker cleaved caspase-3 was detected in neurons and glial cells in the post-mortem brains of influenza patients^46^ and *in vitro* in neurons, astrocytes, and neural stem cells after inoculation with influenza A virus^27,47,48^. Neural activity, which refers to the communication between neural cells, is known to affect learning and memory formation, but this has not been studied in the context of influenza A viruses. Synaptic plasticity, which is the ability to form and strengthen connections between neuronal cells after stimulation, can be studied by investigating neuronal excitability by employing long-term potentiation (LTP) protocols. LTP was reduced in organotypic hippocampal slices of mice inoculated with a mouse-adapted pandemic H3N2 virus in a time-dependent manner^21^, which was associated with a decrease in dendritic spine density^21^. In addition, in mouse-adapted pandemic H1N1 virus inoculated neonatal mice there was a reduction in spontaneous neurotransmission and neuronal excitability^49^. The underlying mechanisms of the changes in neural activity are unclear but might be related to a disbalance in glutamatergic synapse transmission^23^.

Our aim was to gain insight into the neurotropic and neurovirulent potential of recent seasonal and pandemic influenza A viruses. Therefore, we studied the cell tropism, replication kinetics and ability to induce apoptosis of pH1N1 2009 virus, and seasonal H1N1 and H3N2 viruses in a human induced pluripotent stem cell (hiPSC)-derived neural co-culture model that contains neurons and astrocytes. In addition, neural activity and excitability of the neural co-cultures upon virus inoculation were assessed with a micro-electrode array (MEA) platform in order to better understand the underlying mechanisms of neurological complications associated with influenza A virus infection.

## Results

### Inefficient replication of pH1N1 2009, H1N1 2019 and H3N2 2019 viruses in neural co-cultures, consisting of NGN2 neurons and astrocytes

To evaluate the replication efficiency of pH1N1 2009, H3N2 2019 and H1N1 2019 viruses, hiPSC-derived neural co-cultures, consisting of Neurogenin-2 (NGN2) neurons and astrocytes, were inoculated with an multiplicity of infection (MOI) of 1. No increase in infectious virus was observed with any of the viruses up to 10 days post inoculation (dpi; Figure 1A), despite efficient replication in Madin-Darby Canine Kidney (MDCK) cells (Figure S1A). It is possible that inefficient replication occurred in the neural co-cultures inoculated with pH1N1 2009, H1N1 2019 or H3N2 2019 virus as virus titers remained stable and detectable over time, whereas virus titers in culture medium decreased to undetectable levels at 10 dpi (Figure S1B). In addition, intracellular viral RNA levels increased for all three viruses and peaked at 24 hours post inoculation (hpi; Figure 1B).

**Figure 1.**
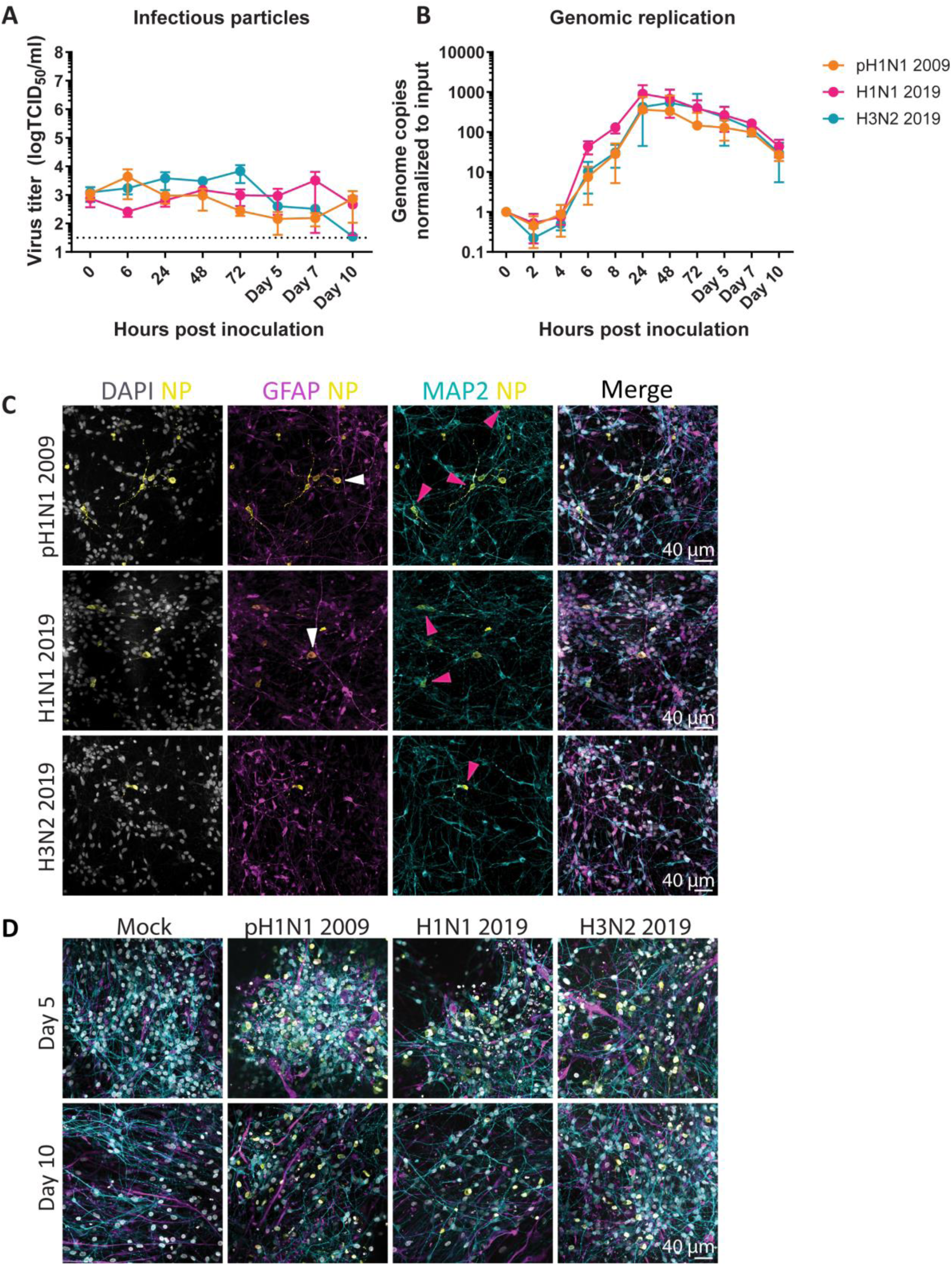
Inefficient replication of pH1N1 2009, H1N1 2019 and H3N2 2019 viruses in neural co-cultures, consisting of NGN2 neurons and astrocytes. Human induced pluripotent stem cell-derived neural co-cultures consisting of NGN2 neurons and astrocytes were inoculated with pH1N1 2009, H1N1 2019 or H3N2 2019 virus with an MOI of 1 at DIV 21. (A) Growth kinetics of pH1N1 2009, H1N1 2019, H3N2 2019. Data represent mean ± standard deviation (SD) and are derived from at least three independent experiments, with three biological replicates and three technical replicates. Dotted line represents lower limited of detection. (B) Intracellular replication kinetics of pH1N1 2009, H1N1 2019 and H3N2 2019. Data represent mean ± SD, and are derived from at least three independent experiments, with one biological replicate and twice in technical duplicates. (C) Neural co-cultures were fixed 72 hours post inoculation and were stained with microtubule-associated protein (MAP2; cyan) as a marker for neurons, glial fibrillary acidic protein (GFAP; magenta) as a marker for astrocytes, and influenza A virus nucleoprotein (NP; yellow) was used to identify infected cells. Cells were counterstained with Hoechst (grey) to visualize the nuclei. Data shown are representative examples from three independent experiments. Co-localization between NP and GFAP is indicated with white arrow, and co-localization between NP and MAP2 with a pink arrow. (D) Neural co-cultures were fixed 5- and 10-days post inoculation and again stained with MAP2, GFAP, NP and DAPI. Data shown are representative examples from three independent experiments.

To investigate the neurotropism, neural co-cultures were fixed at 72 hpi and stained for virus nucleoprotein (NP), microtubule-associated protein (MAP2, neuronal marker) and glial fibrillary acidic protein (GFAP, astrocytic marker). With all viruses, NP^+^ cells were observed in neural co-cultures (Figure 1C). Co-localization of NP was predominantly observed with MAP2 (Figure 1C, pink arrows), although NP occasionally co-localized with GFAP in pH1N1 2009 and H1N1 2019 virus inoculated cultures (Figure 1C, white arrows). In neural co-cultures fixed at 5- and 10 dpi, similar amounts of NP+ cells were detected for all three viruses compared to 3 dpi (Figure 1D).

### No induction of the apoptotic marker cleaved caspase-3 in neural co-cultures inoculated with pH1N1 2009, H1N1 2019 or H3N2 2019 virus

There was no morphological evidence for cell death in the virus inoculated neural co-cultures by visual inspection through light microscope, like cell debris, curling of neural co-cultures or virus-induced cytopathic effects. However, to exclude direct or indirect virus-induced apoptosis in the neural co-cultures, we studied the expression of cleaved caspase-3. Neural co-cultures were fixed at 3-, 5- and 10 dpi and stained for the apoptotic marker cleaved caspase-3 in combination with NP. Cleaved caspase-3 was detected in a few cells in all neural co-cultures, including mock, on day 3, 5 and 10 (Figure 2, Figure S2AB, respectively; indicated with white arrows). There was no clear difference in the number of cleaved caspase-3 expressed cells between mock and virus inoculated neural co-cultures or over time (Figure 2, Figure S2AB). Additionally, we did not observe co-localization of cleaved caspase-3 and NP, suggesting that infected cells did not directly induce apoptosis.

**Figure 2.**
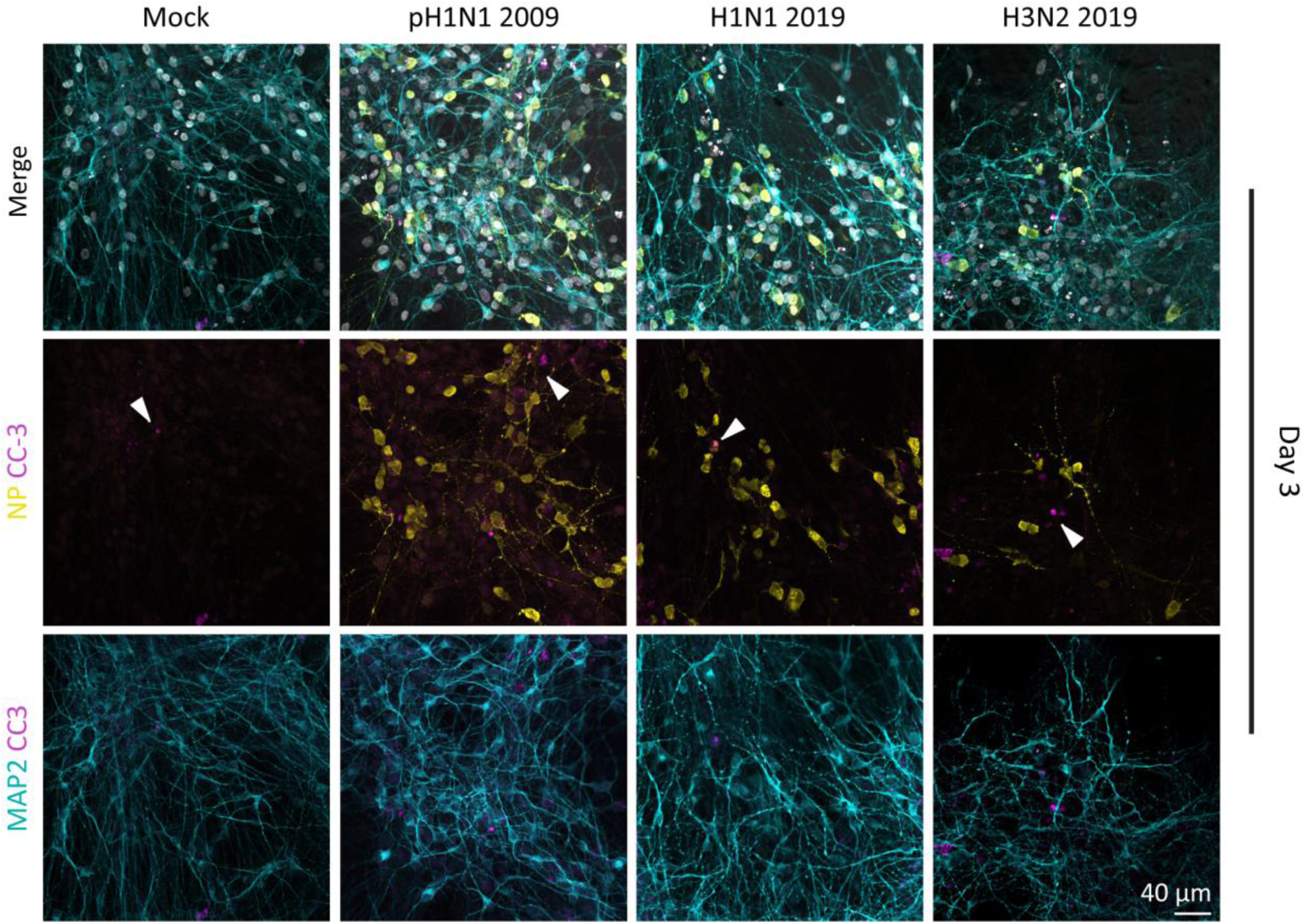
No induction of the apoptotic marker cleaved caspase-3 in the neural co-cultures inoculated with pH1N1 2009, H1N1 2019 or H3N2 2019 virus. Neural co-cultures were fixed 3 days post infection and were stained with microtubule-associated protein (MAP2; cyan) as a marker for neurons, cleaved caspase-3 (CC-3; magenta, indicated with white arrows) as a marker for apoptosis, and influenza A virus nucleoprotein (NP; yellow) to identify infected cells. Cells were counterstained with Hoechst (grey) to visualize the nuclei. Data shown are representative examples from three independent experiments.

### Reduction in spontaneous activity and network activity in neural co-cultures upon inoculation with pH1N1 2009

As we did not observe any differences among the three viruses, we decided to perform all functional analyses on pH1N1 2009 inoculated neural co-cultures that were were fully differentiated at 21 days *in* vitro (DIV) using a MEA system (Axion Systems; MOI 1). The MEA system allows for measuring cell viability and spontaneous activity (Figure 3A). Cell viability and activity can be analyzed by the number of covered and active electrodes. Although the number of covered electrodes did not differ between mock and pH1N1 2009 inoculated wells (Figure S3A), the number of active electrodes was significantly decreased at 5 to 8 and 10 dpi (Figure S3B). This suggests that even though the neural cultures are not dying, their activity is affected. Next, we studied the functionality of the neural co-cultures by measuring the spontaneous activity every 24 hours. Specifically, we analyzed the firing rate (spikes per time interval), inter-spike-interval (ISI) coefficient of variation (measure of spike regularity), burst frequency (bursts per time interval), burst duration, ISI within burst (measure for burst intensity), inter-burst-interval (IBI) coefficient of variation (measure of burst regularity), number of spikes per burst, network burst frequency (network bursts per time duration), network burst duration and number of spikes per network burst (Figure 3A). At baseline level before inoculation, there were no differences in the firing rate, burst frequency or network burst frequency between the cultures dedicated for mock or virus inoculation (Figure S3CDE, respectively). After inoculation, analyses of single electrodes showed that the firing rate and ISI coefficient of variation were significantly reduced between 1 to 7 and 9 dpi in pH1N1 2009 inoculated cultures compared to mock (Figure 3B; Two-way ANOVA F (1, 91) = 9.147 and Figure 3C; Two-way ANOVA F (1, 91) = 17.23, respectively). The burst frequency was significantly reduced at 1 dpi in pH1N1 2009 inoculated cultures compared to mock (Figure 3D; Two-way ANOVA F (1, 89) = 7.278), while the burst duration was significantly reduced at 1 to 6, 8 and 10 dpi (Figure 3E; Two-way ANOVA F (1, 89) = 25.97). There were no differences between the inoculated and mock cultures for the mean ISI within a burst (Figure S3F; F (1, 89) = 1.553), the IBI coefficient (Figure S3G; Two-way ANOVA F (1, 87) = 0.9938). While the number of spikes per burst was significantly decreased (Figure S3H; Two-way ANOVA F (1, 89) = 7.917). Analyses of the multiple electrodes’ parameters revealed that the network burst frequency was significantly decreased at 1 and 2 dpi in pH1N1 2009 inoculated cultures compared to mock (Figure 3F; Two-way ANOVA F (1, 86) = 5.593). The network burst duration was significantly reduced at 1 and 2 to 5 dpi in inoculated cultures compared to mock (Figure 3G; Two-way ANOVA F (1, 86) = 7.470), as well as the number of spikes per network burst, which was reduced at 1 to 3 dpi (Figure S3I; Two-way ANOVA F (1, 86) = 7.859). In conclusion, pH1N1 2009 infection reduced several parameters of spontaneous activity in neural co-cultures impairing neural functioning.

**Figure 3.**
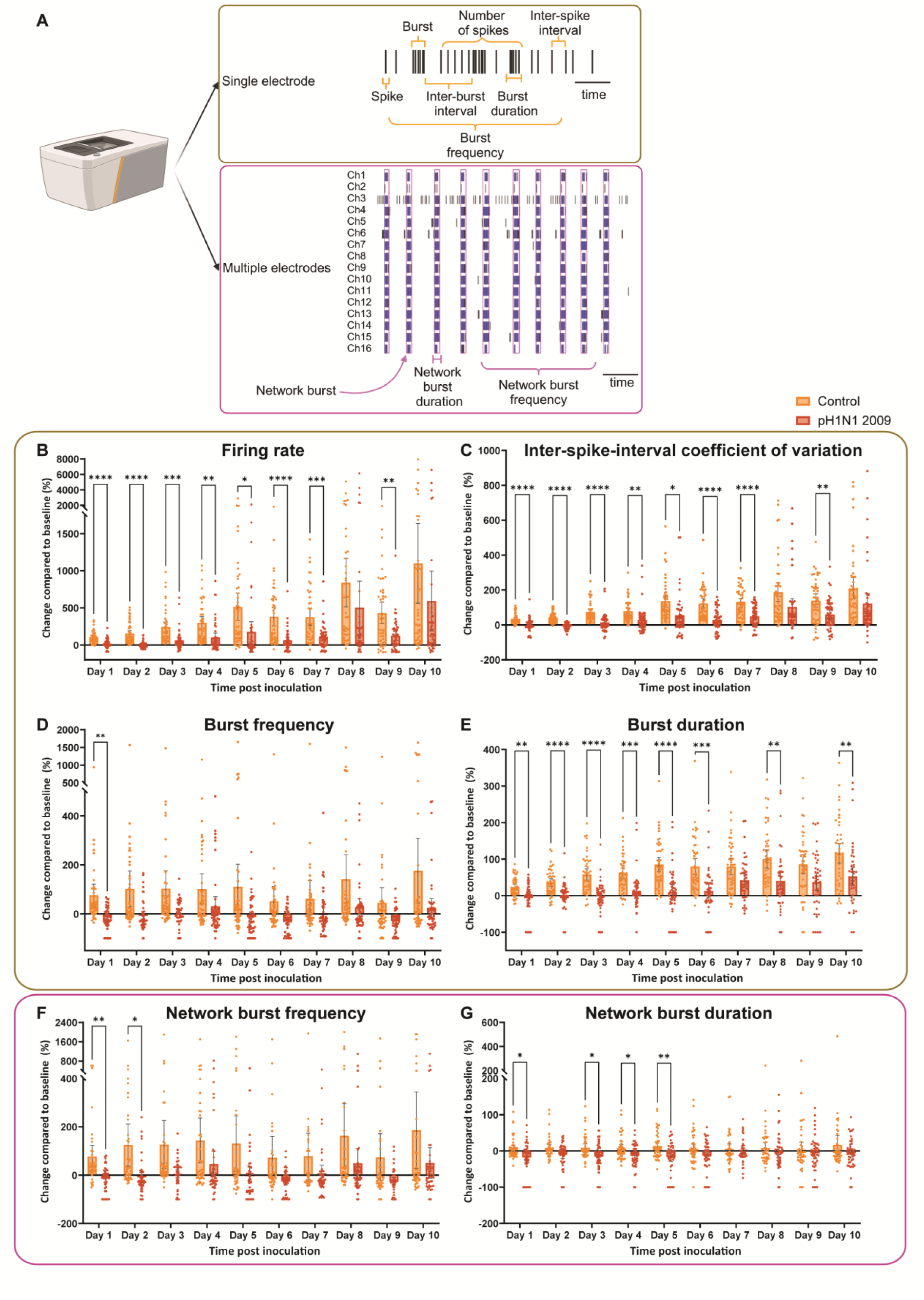
Spontaneous activity measured of neural co-cultures inoculated with pH1N1 2009 virus. Neural co-cultures were mock-treated or inoculated with pH1N1 2009 virus with an MOI of 1 at DIV 21. Spontaneous activity was measured every 24 hours post inoculation for ten days, and recordings were compared to baseline recording obtained before inoculation at DIV 21. The explanation about all parameters measured for spontaneous activity can be found in (A). Created in BioRender. 2, V. (2025) https://BioRender.com/m74f1fe. The following variables were displayed: (B) firing rate, (C) inter-spike-interval coefficient of variation, (D) burst frequency, (E) burst duration, (F) network burst frequency, (G) number of spikes per network burst. Statistical significance was calculated with a two-way analysis of variance (ANOVA) with a Šídák’s multiple comparisons *posthoc* test. Data is displayed from eight independent experiments (n = 48 per group, unless datapoints were excluded based on exclusion criteria see material and methods). Asterisks indicate statistical significance (*P<0.05, **P<0.01, ***P<0.001, ****P<0.0001).

### Number of synapses increases in pH1N1 2009 inoculated neural co-cultures early during infection

Synaptic connections are a necessity for network activity and are refined by neural activity, where connections can be eliminated, maintained or generated. To investigate if a difference in the number of synaptic connections was associated with the observed decrease in neural activity upon inoculation with pH1N1 2009 (Figure 3), the number of synapses was determined at DIV 22 (1 dpi), DIV 24 (3 dpi) and DIV 28 (7 dpi). Neural co-cultures were inoculated with pH1N1 2009, and fixed at 1-, 3- and 7 dpi for immunofluorescence staining for Synapsin I as pre-synaptic marker and Homer1 as post-synaptic marker. We quantified functional synapses by co-localization of Synapsin I and Homer1 (Figure 4A; examples in Figure 4B). A significant increase in synapses was observed at 1- and 3 dpi in pH1N1 2009 inoculated neural co-cultures compared to mock (Figure 4A; day 1: t(62) = 2.776, p = 0.0073; day 3: t(82) = 2.942, p = 0.0042). The number of synapses did not significantly differ at 7 dpi (Figure 4A; day 7: t(62) = 1.703, p-value = 0.0936). In addition, we investigated mRNA levels of the presynaptic glutamate-transporter 1 (vGlut1) as this is known to be affected by neuroinflammation^23,50^. We did not observe any differences in pH1N1 2009 inoculated cultures over time (Figure 4C).

**Figure 4.**
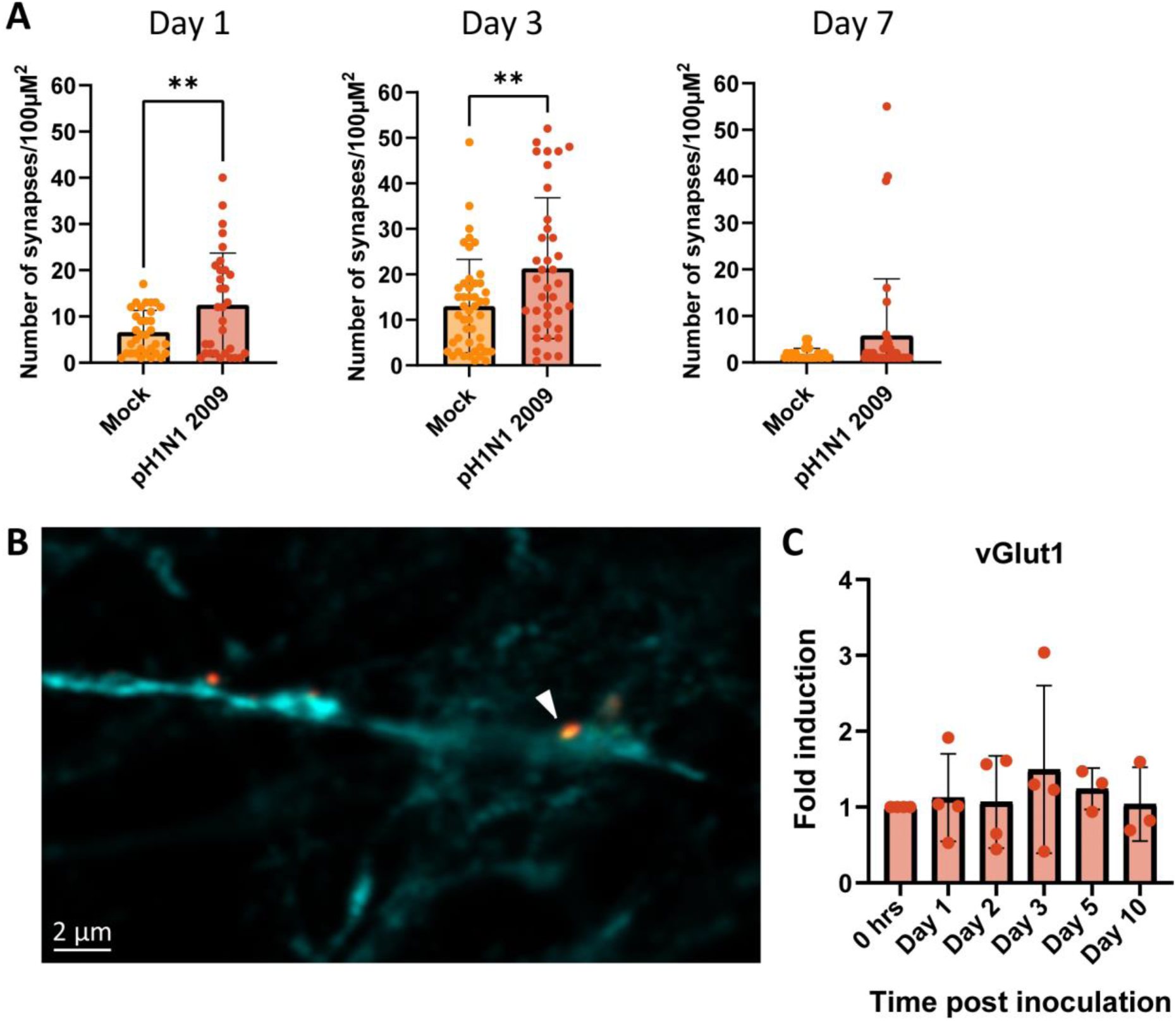
Number of synapses in mock or pH1N1 2009 inoculated cultures. Neural co-cultures were fixed after 1-, 3-, or 7 days post inoculation with pH1N1 2009 virus and stained for Synapsin I as a marker for the presynaptic density (red), Homer1 as a marker for the postsynaptic density (yellow) and with microtubule-associated protein (MAP2; cyan) as a marker for neurons. A synapse was counted when co-localization occurred between Synapsin I and Homer1. (A) Data represent mean ± standard deviation (SD) and are derived from at least two independent experiments, with two technical replicates, where at least 15 images were taken from. Asterisks indicate statistical significance (**P<0.01). (B) An example of a co-localization of Synapsin I and Homer1 is depicted with a white arrow. (C) Expression of vGlut1 in pH1N1 2009 inoculated neural co-cultures over time. Neural co-cultures were inoculated with a MOI of 1 and lysed after 0, 24, 48 or 72 hours and 5- or 10-days post inoculation. Quantitative polymerase chain reaction (qPCR) was performed for β-actin and vGlut1, data is displayed corrected for actin. Data represent mean ± SD, and are derived from at least three independent experiments, with one biological replicate and two in technical replicates.

### Synaptic plasticity is partially impaired after pH1N1 2009 inoculation

Next, we studied synaptic plasticity, which entails the ability to make experience-dependent changes in the strength of neuronal connections, and can be studied by LTP induction. We were able to successfully induce chemical LTP (cLTP) in mock co-cultures at 21 and 31 DIV (Figure S4). At DIV 21, the firing rate was significantly decreased between 0.5 and 24 hours post induction of cLTP (Figure S4A), and only at 0.5 hours post induction of cLTP at DIV 31 (Figure S4B). At both DIV 21 and 31, the burst frequency was significantly increased at all timepoints post induction of cLTP (Figures S4C-D), while the burst duration was significantly reduced at the same timepoints (Figure S4E-F). In addition, the network burst frequency was significantly increased at all timepoints post induction of cLTP (Figure S4G-H), while the network burst duration was significantly reduced at the same timepoints (Figure S4I-J). Next, we inoculated the neural co-cultures with pH1N1 2009 (MOI 1) to further investigate the neurovirulent potential of influenza A viruses on synaptic connections. We induced cLTP at 2 and 10 dpi to observe any acute or post-acute effects. In the pH1N1 2009 virus inoculated cultures the firing rate was significantly decreased at 72 hours post cLTP induction 2 dpi (Figure 5A) and also significantly decreased at 0.5-, 48-, 72- and 96 hours post cLTP induction 10 dpi (Figure 5B). The burst frequency and burst duration did not differ between mock and inoculated cultures at all timepoints post induction of cLTP 2 or 10 dpi (Figure 5C-D-E), with the exception of a significant reduction in the burst duration 4 hours post cLTP induction at 10 dpi (Figure 5F). The network burst frequency did not differ between mock and inoculated cultures post induction of cLTP at 2 dpi (Figure 5G). In the cLTP 10 dpi cultures, the burst frequency was overall lower in the inoculated cultures compared to the mock, which was only significant at 4 hours post induction (Figure 5H). The network burst duration did not alter between mock and inoculated neural co-cultures post induction of cLTP at 2 dpi (Figure 5I), but was significantly decreased at 72-, and 96-hours post induction of cLTP at 10 dpi (Figure 5J). In conclusion, synaptic plasticity is partially impaired later on after inoculation with pH1N1 2009, but not in early stages.

**Figure 5.**
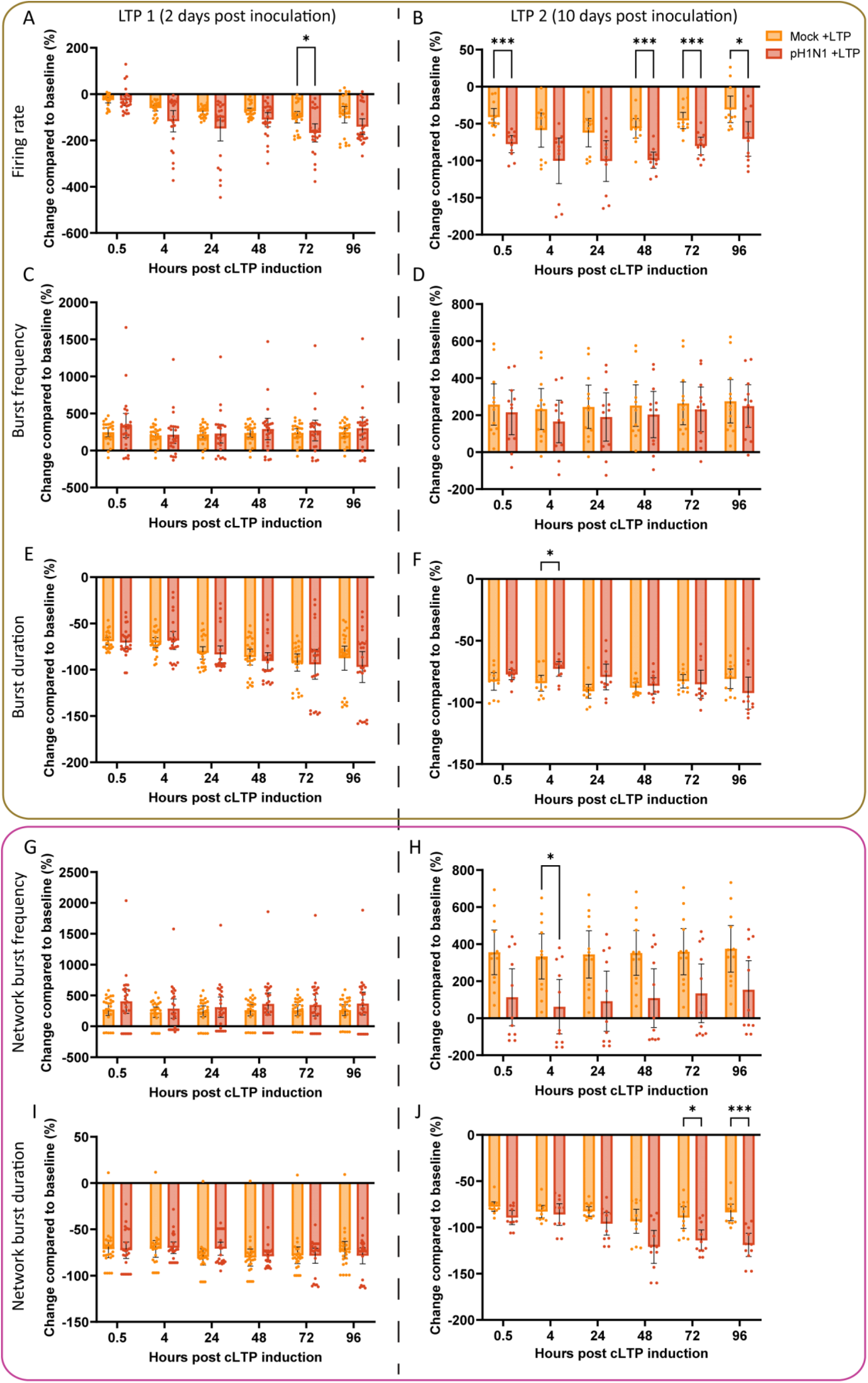
Synaptic plasticity is partially impaired later on after inoculation with pH1N1 2009, but not in early stages. Neural co-cultures were mock-treated or infected with pH1N1 2009 virus with an MOI of 1 at DIV 21. After 2- or 10-days post infection (long-term potentiation 1 (LTP1) and LTP2 respectively), a chemical LTP (cLTP) protocol was employed. Neural activity was measured 0.5-, 4-, 24-, 48-, 72- and 96 hours post induction of cLTP. The following variables were displayed: (AB) Firing rate, (CD) Burst frequency, (EF) Burst duration, (GH) Network burst frequency, (IJ) Network burst duration. Statistical significance was calculated with a two-way analysis of variance (ANOVA) with a Šídák’s multiple comparisons *posthoc* test. Data is displayed from at least four independent experiments (n = 24 per group, unless datapoints were excluded based on exclusion criteria see material and methods) ± 95% CI. Asterisks indicate statistical significance (*P<0.05, **P<0.01, ***P<0.001).

## Discussion

In this study, we revealed that seasonal and pandemic influenza A viruses trigger functional changes in a hiPSC-derived neural co-culture model. Although, replication was inefficient, abundant infection of neurons was observed without the induction of apoptosis. Inoculation of the neural co-cultures with pH1N1 2009 resulted in a reduction of spontaneous neural activity and partially reduced neural excitability.

Overall, the seasonal and pandemic influenza A viruses included in this study did not replicate efficiently or trigger cell death in our neural co-culture model, despite infection of neurons. This aligns with *in vivo* observations, where seasonal or pandemic influenza A viruses are occasionally detected in the CNS. However, these viruses do not seem to replicate efficiently in the CNS in mammals^5,12–20^, like observed for highly pathogenic H5N1 viruses^4^. Histological inflammatory lesions that are associated with active virus replication are not detected in mammals inoculated with seasonal and pandemic influenza A viruses^5,20^. *In vitro* studies have shown that seasonal and pandemic influenza A viruses mostly infected neurons or neuron-like cells^22,25,27^, which is in accordance with our study. However, in a neuroblastoma cell line, pH1N1 2009 virus was able to replicate efficiently^25^, which we did not observe in our hiPSC-derived neural cultures. In general, there are large differences in the replication efficiency between influenza A viruses in neural cells. Highly pathogenic H5N1 viruses replicate more efficiently in cells of the CNS both *in vitro* and *in vivo*^51–53^ compared to seasonal and pandemic influenza A viruses. Sialic acid binding preferences, the multibasic cleavage site or changes in the polymerase genes are suggested to be associated with the differences in neurotropism^4,25^. Since the replication of influenza A viruses is multifactorial, the exact role of viral and host factors that play a role in the neurotropism of influenza A viruses remains to be elucidated

Influenza A virus infection reduces spontaneous neural activity and reduces partially neural excitability in a time-dependent manner. This aligns with a decrease in spontaneous neurotransmission that was observed in mouse-adapted influenza A virus infected mouse hippocampal neurons by whole-cell patch clamping^49^. In our study, neuronal excitability—measured after cLTP stimulation—was only impaired later during infection, but not early during infection. A reduction in neuronal excitability was also observed in mouse-adapted influenza A virus infected mouse hippocampal neurons, which recovered over time^21,49^. In general, there are many mechanisms which can play a role in a reduction of spontaneous neuronal activity or excitability, including cell death, senescence, cytokines and changes in spines or synapses, neurotransmitter release or ion channels. Virus-induced changes can for example be dendritic loss and an impaired glutamate stimulation response, which was observed after Herpes Simplex Virus 1 infection^54^, or altered synaptic activity after SARS-CoV-2 infection^55^. In contrast, neurovirulent cytokines, elicited by a mouse neuroinvasive coronavirus, increased neuronal excitability in mouse neurons^56^. The mechanism by which influenza A virus infection leads to reduced neuronal activity and excitability remains unknown, as it does not appear to involve cell death, synapse loss, or reduced vGlut1 mRNA expression in our study.

Impaired synaptic plasticity is strongly correlated with cognitive impairments, like learning and memory problems. Influenza A virus infection causes cognitive impairments in human cases and experimentally inoculated mice^2,21,39,57^. Impairments in spatial learning and memory, search strategy and cognitive flexibility were observed in H3N2- and H7N2 inoculated mice in a time-dependent manner^21^. H1N1 WSN-inoculated mice showed long-term behavioral changes of anxiety and cognition^57^. The observed reduction of spontaneous neural activity and excitability in pH1N1 2009 inoculated neural co-cultures could play a role in behavior and cognitive impairments as shown by Hosseini *et al*.^21^. The effects of viral infections, like influenza A viruses, on behaviour and cognitive problems is also highlighted in a recent study where non-severe respiratory infections by SARS-CoV-2 and influenza A virus are associated with a similar risk of post-acute sequelae, including neurological symptoms and neurodegenerative disease^42,58^.

We present here a scalable, human-derived model in which we can study the neurotropism and neurovirulent potential of viral infections on a functional level. We show in a human system that –despite the lack of abundant replication and cell death-influenza A virus infection results in neural dysfunction that could contribute, at least in part, to the development of cognitive problems observed in humans. This is important because there is limited knowledge on the neurotropic and neurovirulent potential of a wide range of viruses that are not classically considered as neurotropic or neurovirulent, such as influenza A viruses and SARS-CoV-2. Consequently, the brain is often overlooked while researching these viruses. In addition, so far most *in vitro* studies have used either neuronal cell lines and/or lab-adapted or mouse-adapted viruses, which tend to be more neuropathogenic and therefore do not resemble the neuropathology in human cases. Therefore, this human-derived model is more suitable for studying the neurovirulent potential of human viruses and in the future potential intervention strategies.

Altogether, our findings reveal that pandemic and seasonal influenza A viruses do not replicate efficiently in a hiPSC-derived neural co-culture model. Even though replication is limited, there is neurovirulent potential for recent seasonal and pandemic influenza A viruses in this model, observed by changes in neuronal functioning, like spontaneous neurotransmission and excitability. The insidious changes that we observe in our model can likely contribute to acute and post-acute sequelae of influenza.

## Materials and Methods

### Cells

MDCK (ATCC) cells were cultured in Eagle minimal essential medium (EMEM; Capricorn scientific) supplemented with 10% fetal calf serum (FCS; Sigma), 100 IU/mL penicillin (ThermoFisher Scientific), 100 µg/mL streptomycin (ThermoFisher Scientific), 2 mM glutamine (ThermoFisher Scientific), 1.5 mg/mL sodium bicarbonate (ThermoFisher Scientific), 10 mM HEPES (ThermoFisher Scientific) and 0.1 mM non-essential amino acids (ThermoFisher Scientific). MDCKs were passaged when confluency was over 90%. First, cells were washed with phosphate-buffered saline (PBS) and then dissociated with trypsin-EDTA (0.05%). Medium was refreshed once a week, and cells were kept at 37 °C and 5% CO_2_.

### Human induced pluripotent stem cells

Human induced pluripotent stem cells (hiPSCs; WTC-11 Coriell no. GM25256, provided by Bruce R. Conklin, The Gladstone Institutes and UCSF, San Francisco, CA, USA) were used to generate neural co-cultures. hiPSCs were plated on Matrigel-coated plates (Sigma, 10 µl/mL, resuspended in knockout Dulbecco’s Modified Eagle Medium, KO DMEM; ThermoFisher Scientific) and maintained in hiPSC medium (Table 1). hiPSCs were passaged when confluency of 70-80% was reached. For passaging, hiPSC were washed with PBS and released with Accutase (Life Technologies). Medium was refreshed every other day, and cells were cultured at 37 °C and 5% CO_2_.

**Table 1.**
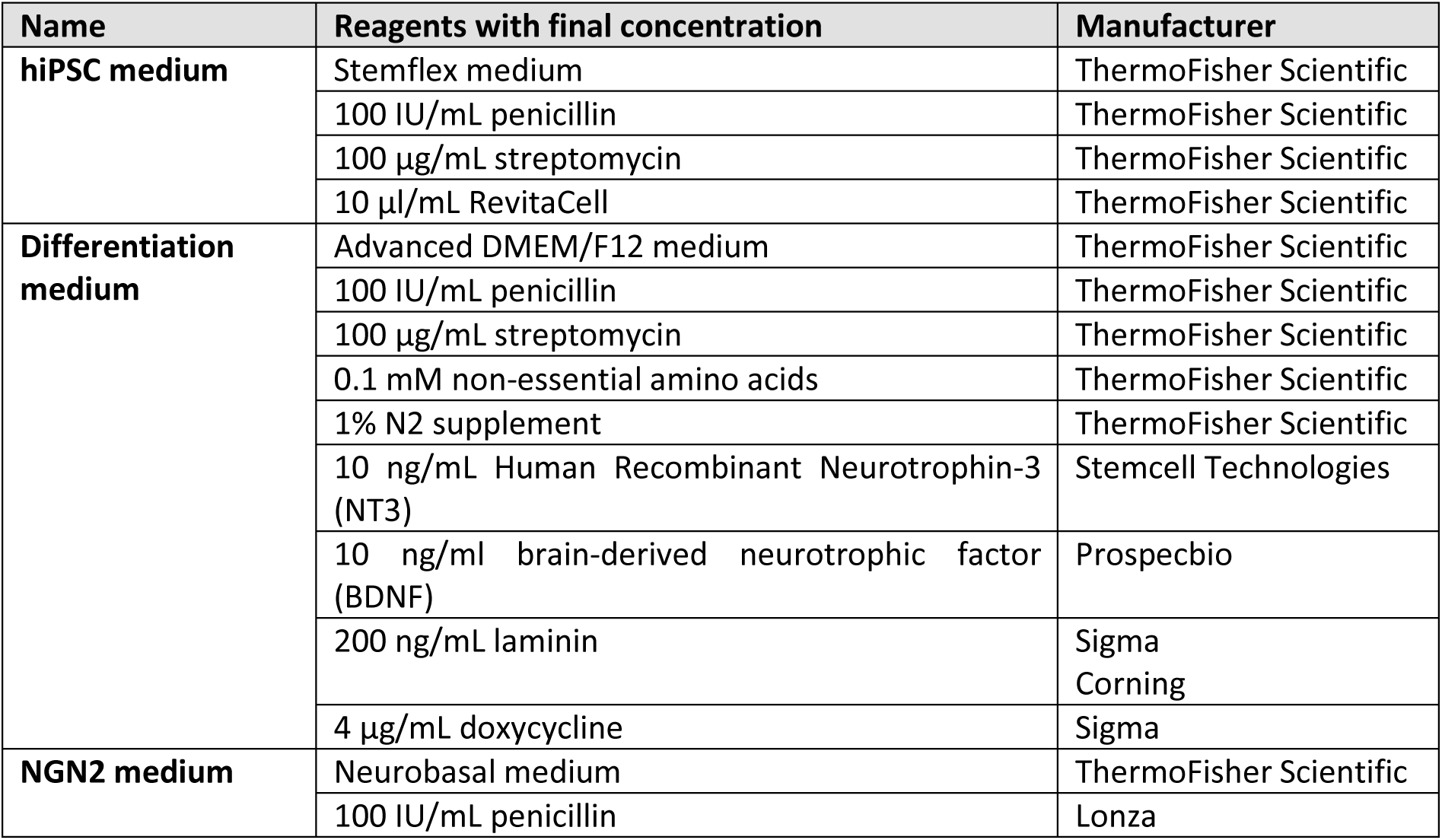
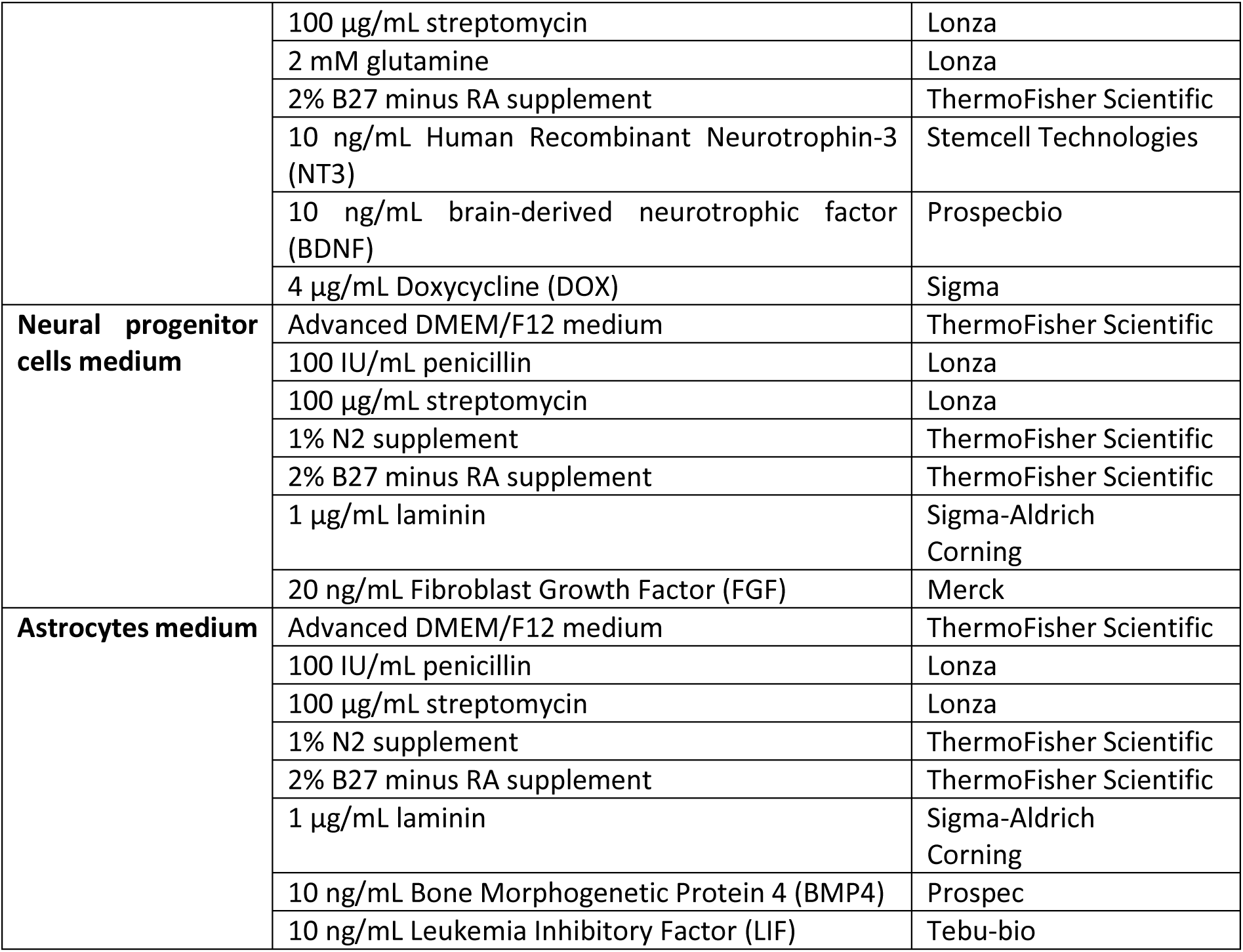
Overview of mediums used for hiPSC-derived NGN2 neurons and astrocytes.

### Differentiation of hiPSCs into NGN2 neuron-astrocyte co-cultures

hiPSCs were differentiated into excitatory cortical neurons by overexpression of neuronal determinant NGN2 via tetracycline-controlled transcriptional activation as previously described^59,60^. In short, coverslips (1.5H, 0117530; Paul Marienfeld) or plates were coated with poly-L-ornithine (Sigma, 100 µg/mL) for 1h at RT in the dark. Then coverslips were washed three times with sterile water and air-dried for 1h. Coverslips or plates were then coated with either laminin (Sigma & Corning, 50 µl/mL, resuspended in DMEM, ThermoFisher Scientific) or Matrigel (Corning, 10 µl/mL resuspended in KO DMEM) for 1h at 37 °C. After incubation, the hiPSCs were plated in hiPSC medium (Table 1) supplemented with 4 µg/mL fresh doxycycline (Sigma). The next day, the medium was refreshed with differentiation medium (Table 1). In every medium, growth factors were added (fresh) before refreshing. In order to guarantee the formation of functional synapses and thus functional synaptic plasticity within the network, hiPSC-derived astrocytes are needed for the NGN2 neurons to mature^59,60^. Therefore, on day 3, hiPSC-derived astrocytes were added to the culture in a 1:1 ratio. hiPSC-derived astrocytes were differentiated as described previously from hiPSC-derived neural progenitor cells (NPCs)^61^. During maintainance, and differentiation of hiPSC-derived NPCs and astrocytes, medium was refreshed with astrocyte or NPC medium (Table 1) every other day. Cells were were split when confluent, roughly once per week. On day 4, medium of the co-cultures was refreshed with NGN2 medium (Table 1). During the differentiation and maturation of neural co-cultures, half of the medium was refreshed every other day. After DIV 21, the neural co-cultures were used in experiments. All cells were kept at 37 °C and 5% CO_2_.

### Viruses

Pandemic 2009 H1N1 virus (pH1N1 2009, A/Netherlands/602/2009) was isolated from a patient who visited Mexico and the virus was propagated twice in MDCK cells. Seasonal H1N1 virus and H3N2 virus isolated from 2019 were kindly provided by the National influenza centrum (Table 2) and the viruses were propagated once in MDCK cells.

**Table 2.** Subtype and clade of 2019 viruses and 2009 virus.

| **Virus** | **Subtype** | **Clade** | **Dutch identification number** |
| --- | --- | --- | --- |
| H3N2 2019 | AH3N2 | 3C.3a | 19A01618 |
| H1N1 2019 | AH1N1pdm09 | 6B.1A5A | 19A01659 |
| H1N1 2009 | AH1N1pdm09 | 6B.1 | GISAID Isolate ID: EPI_ISL_31217 |

### Virus infection

Before infection, medium was removed and cells were inoculated with pH1N1 2009, H1N1 2019 or H3N2 2019 viruses at the indicated MOI. Viruses were diluted in NGN2 medium (Table 1) for neural co-cultures, and in infection medium for MDCKs (EMEM supplemented with 100 IU/mL penicillin, 100 µg/mL streptomycin, 2 mM glutamine, 1.5 mg/mL sodium bicarbonate, 10 mM HEPES, 0.1 mM nonessential amino acids, and 1 µg/µL TPCK-treated trypsin (Sigma)). After 1h of infection MDCK cells were washed three times to remove residual unbound virus. For infection of the hiPSC-derived neural co-cultures, virus-containing medium was replaced with fresh NGN2 medium (Table 1), without washing. For infection of hiPSC-derived neural co-cultures on micro-electrode array (MEA) plates, virus-containing medium was not removed, fresh medium was added with the virus-containing medium 1:1.

### Virus titration

Virus titers were determined by endpoint dilution on MDCK cell as described before^62^. In short, 10-fold serial dilutions in triplicates of cell supernatant were prepared in infection medium for MDCKs. MDCKs were washed once with plain EMEM to remove residual FCS prior to adding the diluted supernatants to the MDCKs. After 1h incubation period at 37°C and 5% CO_2_, inoculum was removed and fresh infection medium was added to the cells. The supernatants of the infected MDCKs were tested for agglutination three days after inoculation. For that, supernatant was mixed with 0.33% turkey red blood cells and incubated for one hour at 4°C. The titers of infectious virus were calculated according to the method of Spearman and Kärber^63,64^, and expressed as TCID_50_/mL. All experiment with infectious pH1N1 2009, H1N1 2019 and H3N2 virus were performed in a class II biosafety cabinet under biosafety level 2 (BSL-2) conditions at the Erasmus Medical Center.

### RNA isolation, qRT-PCT, cDNA production and qPCR

Influenza A virus RNA was detected in samples using a quantitative reverse-transcription polymerase chain reaction (qRT-PCR) assay targeting the M gene segment as described previously^65^. In short, RNA was isolated by using a MagnaPure LC system with a MagnaPure LC total nucleic acid isolation kit (Roche Diagnostics). The concentration of RNA was determined using a NanoDrop spectrophotometer. Influenza A virus was detected by targeting the matrix gene (M; table 3). The PCR program was as follows: (2 min 50°C, 20 sec 95°C followed by 45 cycli of 3 sec 95°C and 30 sec 60°C). Reactions were performed on a 7500 Real-Time PCR system (Applied Biosystems).

**Table 3.**
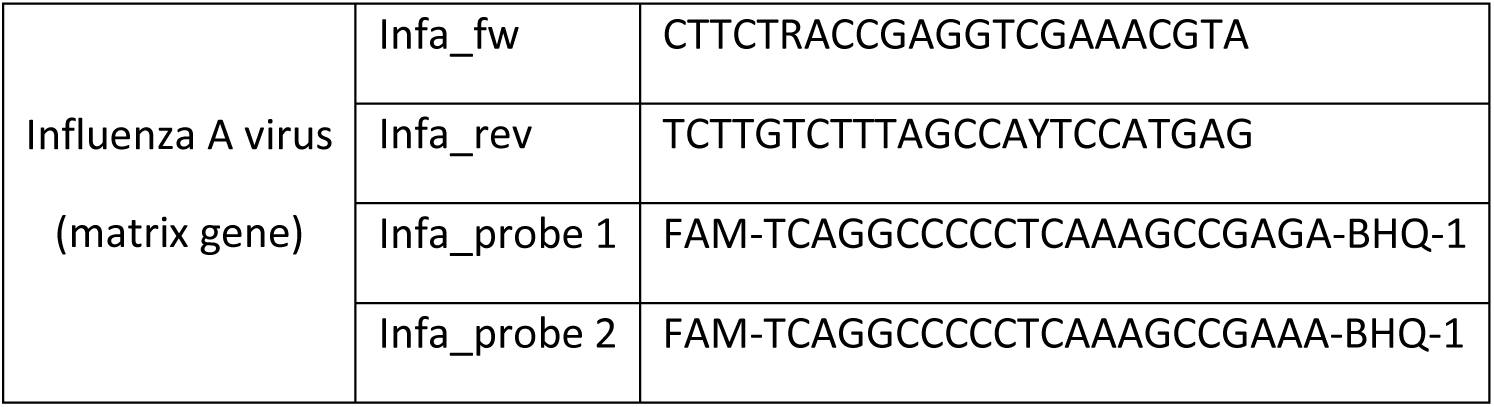
Primer-probes targeting the matrix gene of influenza A virus.

500 ng RNA was reverse transcribed into cDNA using the SuperScript IV reverse transcriptase (Invitrogen) according to the manufacturer’s protocol. Subsequently, the presence of vGlut1 was evaluated by qPCR (table 4), on a 7500 Real-Time PCR system (Applied Biosystems). Fold changes were calculated using the 2^-ΔΔCt^ method. Normalization was performed using the mean Ct values of household gene Actin as a loading control for every sample.

**Table 4.**
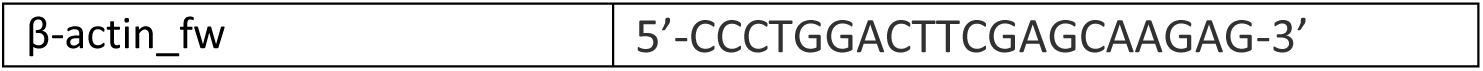
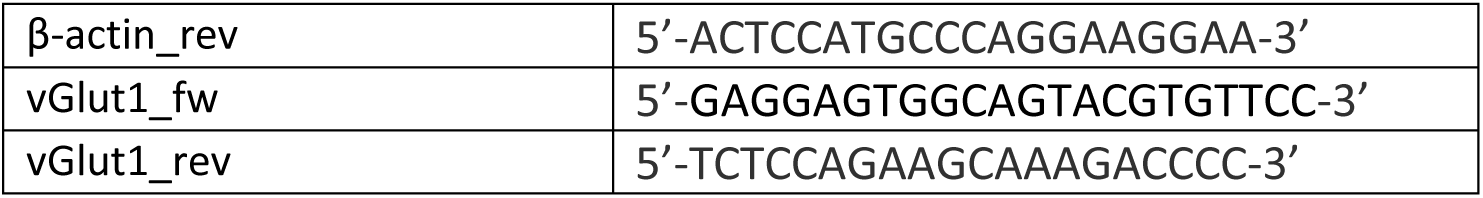
Gene-specific primers for vGlut1 and Actin.

### Immunofluorescence staining

Cells were fixed using 10% formalin for 30 minutes and washed afterwards with PBS. Cells were permeabilised for 15 min using 1% Triton X-100 in PBS and blocked with in PBS supplemented 0.5% Triton X-100 and 1% bovine serum albumin for 30 min at RT. Primary antibodies (Table 5) incubation was performed for 1h at RT in blocking solution, except for α-Homer1, α-Synapsin I, which were incubated O/N at 4°C. After washing with blocking solution, secondary antibodies conjugated to Alexa-488, Alexa-555, Alexa-647 (Invitrogen, Table 5) were used with the corresponding primary antibody and incubated for 1h at RT in blocking solution. Slides were washed in blocking solution and incubated with a solution of Hoechst (ThermoFisher Scientific) in blocking buffer for 15 min. Slides were again washed in blocking solution, dipped in water and mounted in ProLong Glass Antifade Mountant (ThermoFisher Scientific). Samples were imaged using a Zeiss LSM 700 confocal microscope and synapses were images using a Zeiss LSM 900 confocal microscope.

**Table 5.**
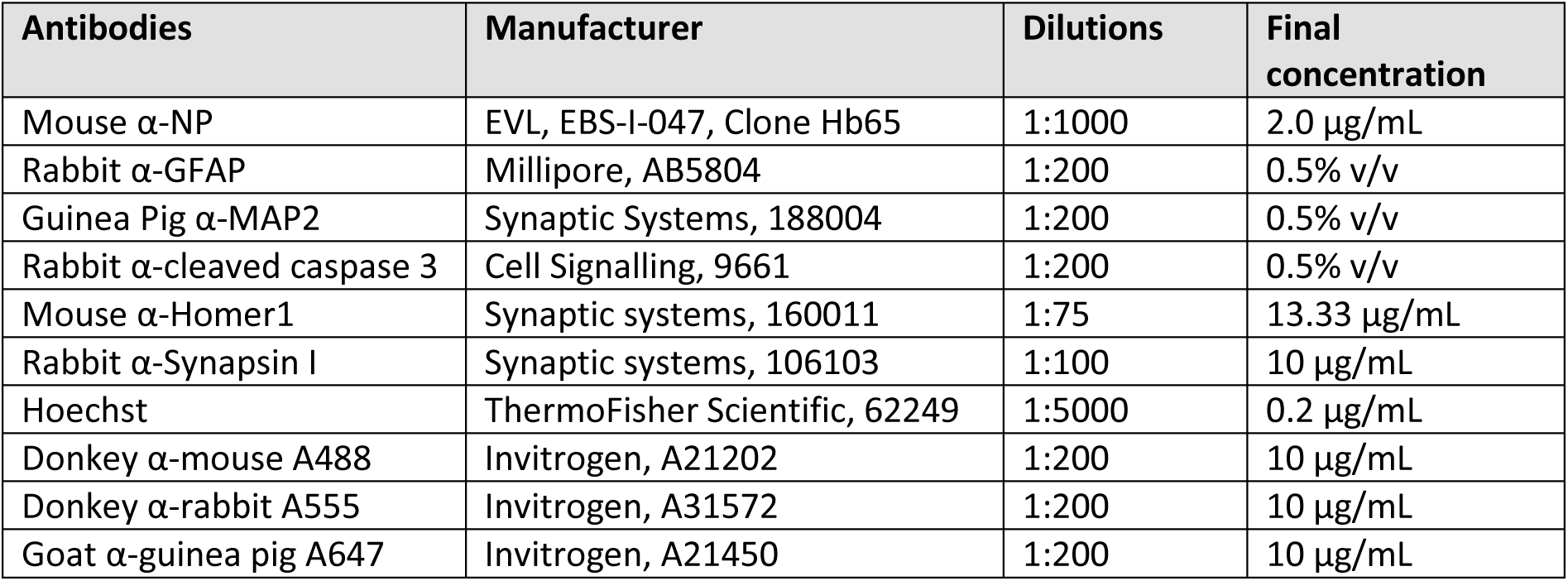
List of used antibodies and according dilutions.

### Microscopic analysis for synapses

Neural co-cultures were mock-inoculated or inoculated with pH1N1 2009 as described earlier and fixed after 24-, 72 hpi or 7 dpi. Neural co-cultures were stained for Homer1 and Synapsin I as described earlier to quantify the number of synapses. Confocal images were taken with a Zeiss LSM 900 module on an Axio Imager Z2 upright microscope fitted with a 63× 1.4NA plan-apochromat oil objective. For imaging the Hoechst, Alexa Fluor 488, Alexa Fluor 555 and Alexa Fluor 647 dies, samples were excited using a 405nm, 488nm, 561nm and 640nm diode laser respectively. Emission was filtered appropriately for each die with 400-600 nm, 410-546 nm (SP545), 540-617 nm, 645-700 nm filter ranges, respectively. ZenBlue image acquisition software (Carl Zeiss gmbh Oberkochen) was used to capture all images. Pixelsize was set to 319.5 µm x 319.5 µm for optimal resolution (1024 x 1024 pixels). Per timepoint at least 15 images were taken per coverslip from two independent experiments. Using of FIJI (version 1.54f), puncta of Synapsin I and Homer1 were selected by the function Find Maxima. A synapse was counted when co-localization occurred between Synapsin I and Homer1. We used a FIJI plugin (‘Nearest Neighbours’) available at https://github.com/ErasmusOIC/NearestNeighbourAnalysis to determine the distance between each Homer1 puncta and its nearest Synapsin I neighbour. Subsequently we defined puncta as co-localizing if the distance between the Homer1 and Synapsin I was below 250 nm.

### MEA recordings of spontaneous neural activity in NGN2 neurons co-cultured with astrocytes

For MEA recordings, hiPSC-derived neural co-cultures were plated onto 24-well MEA plates (Axion Biosystems) as described before. Plates were acclimatized for 5 min before spontaneous activity was recorded for 3 min every 24 hours using the Axion Biosystems Maestro MEA at 37°C and 5% CO_2_. Data analysis was performed using AxIs software (Axion Biosystems Inc.). Covered electrodes were defined as electrodes with a minimum resistance of 18 kΩ. We excluded wells for further analysis if ≤5 electrodes were covered. Active electrodes were defined as electrodes with a minimum of five spikes per minute. We excluded wells for analysis if a well had ≤5 active electrodes at the baseline recording. The threshold for spike detection was defined as ≥6-fold the standard deviation (SD) of the root mean square noise. The threshold for burst detection was defined as >5 spikes within a time window of 100 ms. The threshold for network burst detection was defined as >50 spikes within a time window of 100 ms, with a minimum of 50% participating electrodes per well.

### Induction of chemical long-term potentiation in NGN2 neurons co-cultured with astrocytes

We employed a cLTP protocol demonstrated earlier in human iPSC-derived neuronal network cultures^66^ to study synaptic plasticity in our NGN2 neurons co-cultured with astrocytes. In short, forskolin (Tocris; final concentration 50 μM) and rolipram (Tocris; final concentration 0.1 μM) were dissolved in DMSO and further diluted in NGN2 medium. A maximum amount of 50 µL diluted drug mixture were added to the well, not exceeding a total of 500 µL (Table 1). As negative control, the same amount of DMSO was diluted in NGN2 medium (Table 1). These mixtures were pre-warmed for at least 15 minutes at 37°C. A baseline recording was performed before inducing cLTP. The drug mixtures were added directly in the medium, and neural activity was measured after 30 minutes incubation. After incubation, drugs were washed out as follows: 100 µl of fresh medium per well was removed and 100 µl of fresh medium per well was added, this was repeated three times. Then 150 µl of fresh medium per well was removed and 150 µl of fresh medium per well was added, this was repeated two times. After wash-out, we recorded neural activity after 4, 24, 48, 72 and 96 hours. Settings for MEA recordings and analysis were done exactly the same as for the spontaneous neural activity recordings. Data was normalized based on the average values of the negative control wells per experiment.

### Statistical analysis

Statistical differences between experimental groups were determined as described in the figure legends. P values of ≤ 0.05 were considered significant. Graphs and statistical tests were made with GraphPad Prism version 10.4.2. Figures were prepared with ImageJ 1.54p and Adobe Illustrator 27.8.1.

## Acknowledgements

We thank Syriam Sooksawasdi Na Ayudhya and Eleanor Marshall for technical support. We also thank Dr. Johny Pires and colleagues of Axion BioSystems for useful discussions and technical support. This work was funded in part by a fellowship from the Netherlands Organization for Scientific Research (VIDI contract 91718308) and in part funded by the European Union under grant agreement (101084171) - (Kappa-Flu). Views and opinions expressed are however those of the author(s) only and do not necessarily reflect those of the European Union or REA. Neither the European Union nor the granting authority can be held responsible for them. L.B. is supported by The Netherlands Organization for Scientific Research (XS contract number OCENW.XS22.2.045) and by a grant 2023 from the European Society of Clinical Microbiology and Infectious Diseases (Europäische Gesellschaft für klinische Mikrobiologie und Infektionskrankheiten) (ESCMID). B.L., S.K and F.V are supported by the Netherlands Organ-on-Chip Initiative, an NWO Gravitation project (024.003.001) funded by the Ministry of Education, Culture and Science of the government of the Netherlands.

## Author Contributions

**Conceptualization:** Feline F. W. Benavides, Lisa Bauer, Debby van Riel

**Data Curation:** Feline F. W. Benavides, Lisa Bauer

**Formal analysis:** Feline F. W. Benavides, Lisa Bauer, Johan A. Slotman, Annabel L. V. Kempff

**Funding Acquisition:** Debby van Riel, Lisa Bauer

**Investigation:** Feline F. W. Benavides, Lisa Bauer, Marla Lavrijsen, Annabel L. V. Kempff

**Methodology:** Feline F. W. Benavides, Lisa Bauer, Hilde Smeenk, Bas Lendemeijer, Marla Lavrijsen, Johan A. Slotman, Annabel L. V. Kempff

**Project Administration:** Feline F. W. Benavides, Lisa Bauer

**Resources:** Lisa Bauer, Femke M.S. de Vrij, Steven A. Kushner, Johan A. Slotman

**Software:** Feline F. W. Benavides, Johan A. Slotman

**Supervision**: Femke M.S. de Vrij, Steven A. Kushner, Lisa Bauer, Debby van Riel

**Validation:** Feline F. W. Benavides, Lisa Bauer, Annabel L. V. Kempff

**Visualization:** Feline F. W. Benavides, Lisa Bauer, Annabel L. V. Kempff

**Writing – Original Draft Preparation:** Feline F. W. Benavides, Lisa Bauer, Debby van Riel

**Writing – Review & Editing:** all authors

