## Supplemental information for "Influenza A Virus Infection Impairs Neuronal Activity in Human iPSC-Derived NGN2 Neural Co-Cultures"

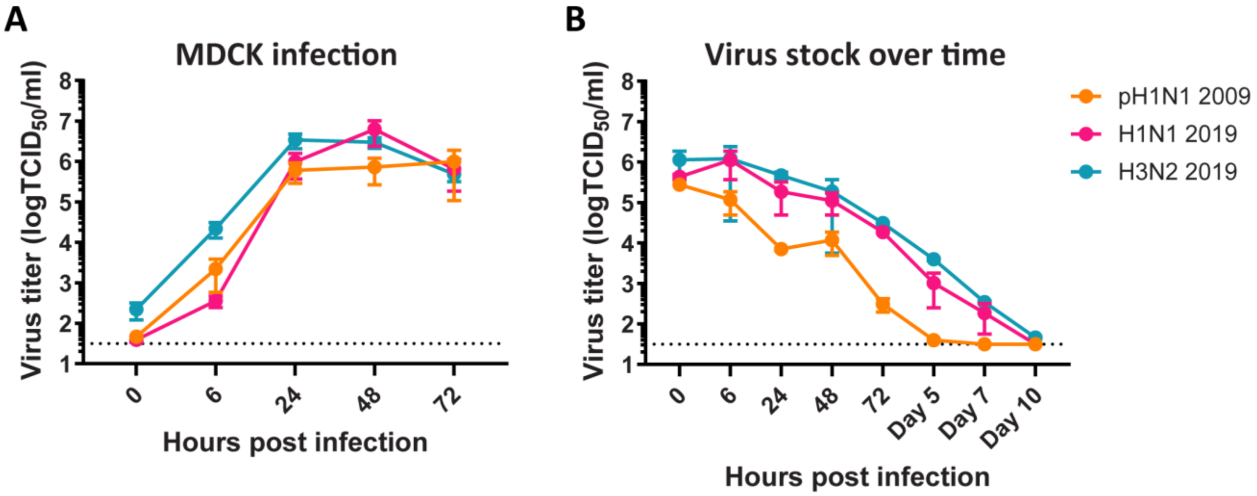


**Fig S1. Replication of pH1N1 2009, H3N2 2019 and H1N1 2019 virus in MDCK cells and without cells.** (A) Madin-Darby Canine Kidney (MDCK) cells were inoculated in parallel with neural co-cultures with pH1N1 2009, H1N1 2019 and H3N2 2019 with a MOI of 1. (B) Virus stocks of pH1N1 2009, H1N1 2019 and H3N2 2019 virus were monitored over time without cells to study the stability of the viruses over time.

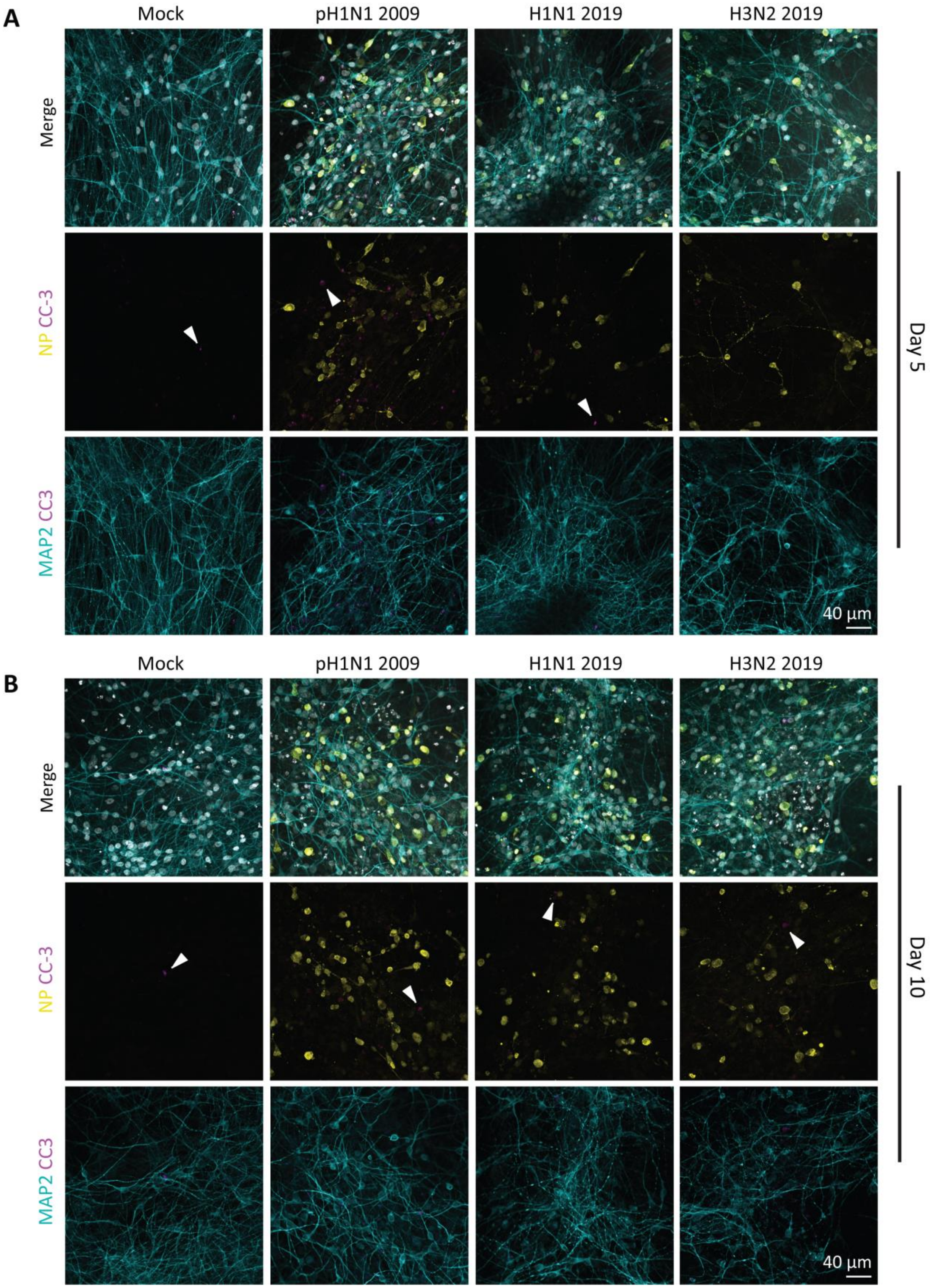
A
Mock
pH1N1 2009
H1N1 2019
H3N2 2019
Merge
NP CC-3
MAP2 CC3
Day 5
40 µm
B
Mock
pH1N1 2009
H1N1 2019
H3N2 2019
Merge
NP CC-3
MAP2 CC3
Day 10
40 µm

**Figure S2. No induction of the apoptotic marker cleaved caspase-3 in the neural co-cultures inoculated with pH1N1 2009, H1N1 2019 or H3N2 2019 virus.** Neural co-cultures were fixed (A) 5 days post infection (dpi) or (B) 10 dpi and were stained with microtubule-associated protein (MAP2; cyan) as a marker for neurons, cleaved caspase-3 (CC-3; magenta, indicated with white arrows) as a marker for apoptosis, and influenza A virus nucleoprotein (NP; yellow) to identify infected cells. Cells were counterstained with Hoechst (grey) to visualize the nuclei. Data shown are representative examples from three independent experiments.

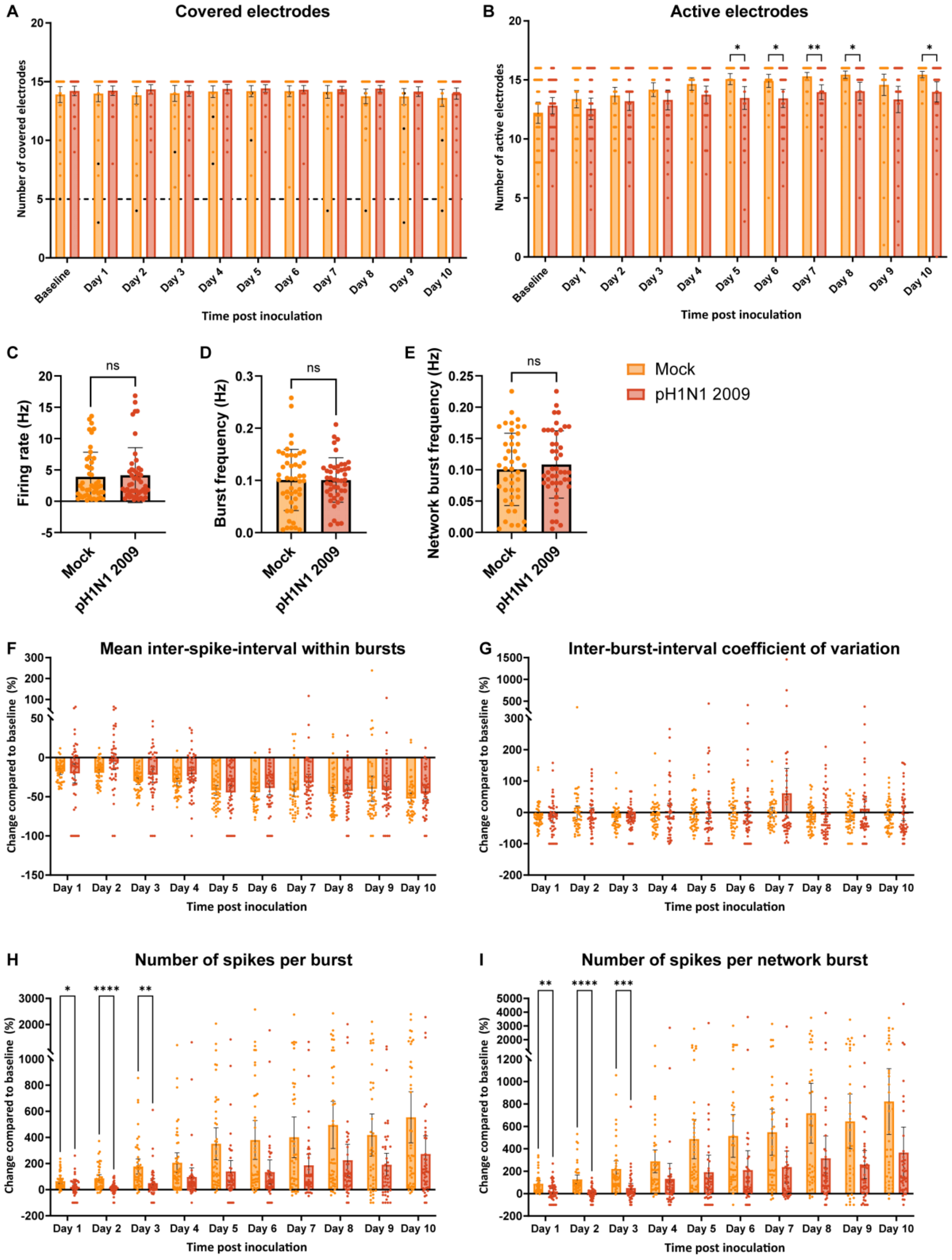


**Figure S3. Spontaneous activity measured of neural co-cultures infected with pH1N1 2009 virus.** Neural co-cultures were mock-treated or inoculated with pH1N1 2009 virus with an MOI of 1 at DIV 21. Spontaneous activity was measured every 24 hours post inoculation for ten days, and recordings were compared to baseline recording obtained before infection. As measures for cell viability, the variables (A) covered electrodes and (B) active electrodes are displayed. At baseline, the firing rate (C), burst frequency (D) and network burst frequency (E) were compared between designated mock and inoculated groups. Other variables for spontaneous activity that were displayed: (F) mean inter-spike-interval within bursts, (G) inter-burst-interval coefficient of variation, (H) number of spikes per burst and (I) number of spikes per network burst. Statistical significance was calculated with a two-way analysis of variance (ANOVA) with a Šídák's multiple comparisons *posthoc* test for (AB) and (FGHI). Statistical significance was calculated with a Welch’s t test for (CDE). Data is displayed from eight independent experiments (n = 48 per group, unless datapoints were excluded based on exclusion criteria see material and methods). Asterisks indicate statistical significance (*P<0.05, **P<0.01, ***P<0.001, ****P<0.0001).

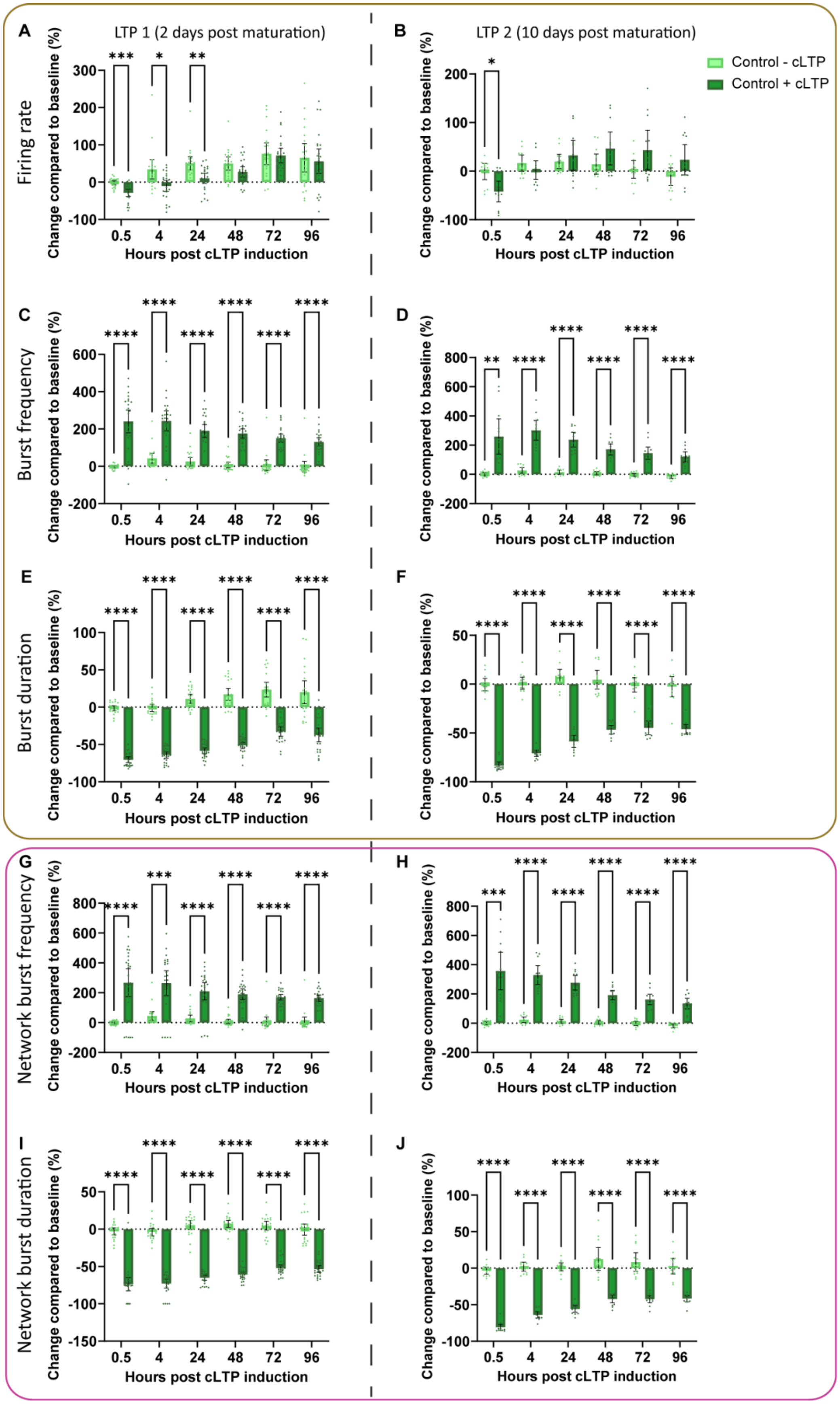

LTP 1 (2 days post maturation)
LTP 2 (10 days post maturation)
Control - cLTP
Control + cLTP
Firing rate
Burst frequency
Burst duration
Network burst frequency
Network burst duration
Change compared to baseline (%)
Hours post cLTP induction

**Figure S4. Chemical long-term potentiation is successful induced in neural co-cultures at DIV 23 and 31.** Neural co-cultures were treated with forskolin and rolipram to induce chemical long-term potentiation (cLTP) at 2- or 10 days post differentiation (+cLTP), or treated with DMSO as control (-cLTP). Neural activity was measured 0.5-, 4-, 24-, 48-, 72- and 96 hours post cLTP induction. The following variables were displayed: (AB) Firing rate, (CD) Burst frequency, (EF) Burst duration, (GH) Network burst frequency, (IJ) Network burst duration. Statistical significance was calculated with a two-way analysis of variance (ANOVA) with a Šídák's multiple comparisons *posthoc* test. Data is displayed from at least four independent experiments (n = 24 per group, unless datapoints were excluded based on exclusion criteria see material and methods) ± 95% CI. Asterisks indicate statistical significance (*P<0.05, **P<0.01, ***P<0.001, ****P<0.0001).
